# Do male and female heterogamety really differ in expression regulation? Lack of global dosage balance in pygopodid geckos

**DOI:** 10.1101/2020.06.03.132241

**Authors:** Michail Rovatsos, Tony Gamble, Stuart V. Nielsen, Arthur Georges, Tariq Ezaz, Lukáš Kratochvíl

## Abstract

Differentiation of sex chromosomes is thought to have evolved with cessation of recombination and subsequent loss of genes from the degenerated partner (Y and W) of sex chromosomes, which in turn leads to imbalance of gene dosage between sexes. Based on work with traditional model species, theory suggests that unequal gene copy numbers lead to the evolution of mechanisms to counter this imbalance. Dosage compensation, or at least achieving dosage balance in expression of sex-linked genes between sexes, has largely been documented in lineages with male heterogamety (XX/XY sex determination), while ZZ/ZW systems are assumed to be usually associated with the lack of chromosome-wide gene dose regulatory mechanisms. Here we document that although the pygopodid geckos evolved male heterogamety with a degenerated Y chromosome 32-72 million years ago, one species in particular, Burton’s legless lizard (*Lialis burtonis*), does not possess dosage balance in the expression of genes in its X-specific region. We summarize studies on gene dose regulatory mechanisms in animals and conclude that there is in them no significant dichotomy between male and female heterogamety. We speculate that gene dose regulatory mechanisms are likely to be related to the general mechanisms of sex determination instead of type of heterogamety.

## Introduction

Differentiated sex chromosomes evolved independently in numerous animal and plant lineages^1^. The differentiation is connected with cessation of recombination and subsequent loss of functional genes from the Y or W sex chromosomes which leads to gene dose differences between sexes. Selection will favour the evolution of mechanisms that regulate these differences at the cellular level, as alterations in gene copy number generally alters gene expression, ultimately impacting cell physiology and organismal fitness^2–5^. Different taxa have evolved distinct strategies to regulate the unequal gene copy numbers and the associated gene dosage imbalances between the sexes related to differentiated sex chromosomes^6^. The most well-known mechanism is dosage compensation, which restores the expression of X or Z-specific genes in the heterogametic sex to the ancestral expression levels^7–9^. Dosage compensation usually leads to dosage balance, i.e. equal expression levels of the X or Z-specific genes between the sexes, however some animal lineages can reach dosage balance in the expression between sexes without keeping the ancestral expression level. Other animal lineages do not compensate and balance expression in the majority of the sex-linked genes at either the level of transcription or translation^10,11^. Dosage compensation or at least dosage balance between sexes was documented largely in lineages with male heterogamety (XX/XY sex determination) such as in several insect lineages, nematode worms, the green anole and eutherian mammals, with sticklebacks, basilisks and platypus being exceptions^6,12,13^. On the contrary, ZZ/ZW systems are usually associated with the lack of chromosome-wide gene dose regulatory mechanisms, often referred to as “partial” or “incomplete” dosage compensation. In such cases, it is assumed that the epigenetic mechanisms regulating gene expression in the heterogametic sex are restricted to a few dosage sensitive genes on the Z chromosome where changes in gene dosage are tied to deleterious fitness effects or lethality, whereas the majority of the genes display different expression levels in males and females^6,14^. This implies that some genes are dosage sensitive (low heterozygote fitness or lethality) whereas others are less so. The lack of chromosome-wide dosage compensation and dosage balance has been documented in parasitic blood flukes, tonguefish, caenophidian snakes, birds, a trionychid turtle and the Komodo dragon, with lepidopteran insects and *Artemia franciscana* representing the only known exceptions here^6,11,15,16–18 and unpublished manuscript^.

It is assumed that a dichotomy in the gene dose regulatory mechanisms between male and female heterogamety occurs, and several, mostly adaptive explanations have been suggested to explain this pattern^19–24^. The hypothesis of differences in gene dose regulation mechanisms between male and female heterogamety is supported from studies of a limited number of lineages across animals (i.e. mainly nematodes, insects, vertebrates), with notably different embryonic (and mainly gonadal) development, highly dissimilar sex chromosome gene content, and genome organization. We argue that this conclusion was premature. To study this phenomenon effectively, we need to explore patterns within a single, phylogenetically coherent lineage with variable sex determining modes. Amniotes (mammals and sauropsids) evolved sex chromosomes independently around 40 times, with geckos representing about half of the recorded transitions^25,26^. Currently, we know genes linked to sex chromosomes in only 16 amniote lineages with putative independently evolved sex chromosomes (reviewed in ^13,27^) and gene dose regulatory mechanisms were studied in just eight of these lineages (Table 1). In our quest for understanding the evolution of sex determination and gene dose regulatory mechanisms, we focus here on the pygypodid geckos (family Pygopodidae).

**Table 1:**
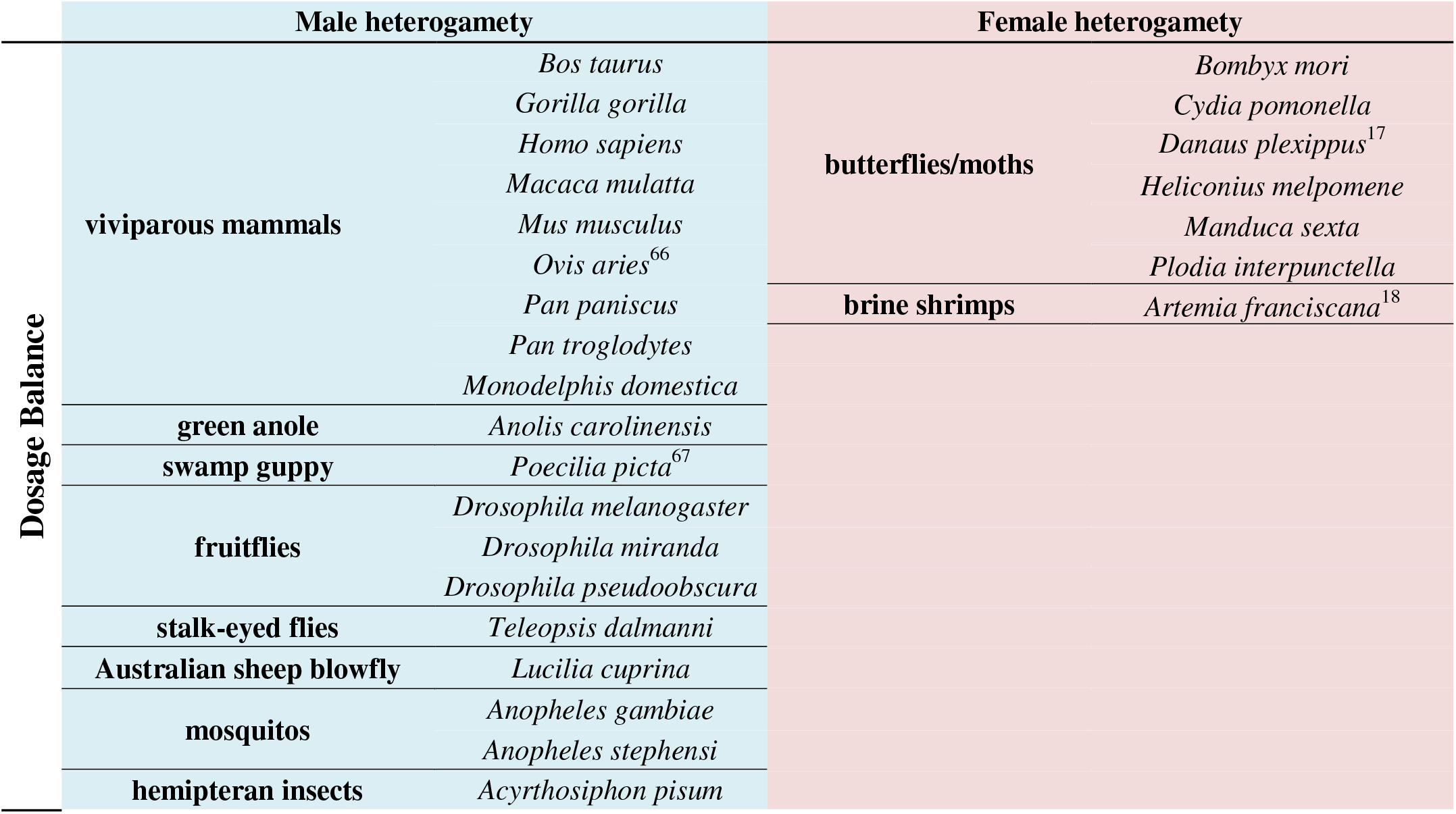
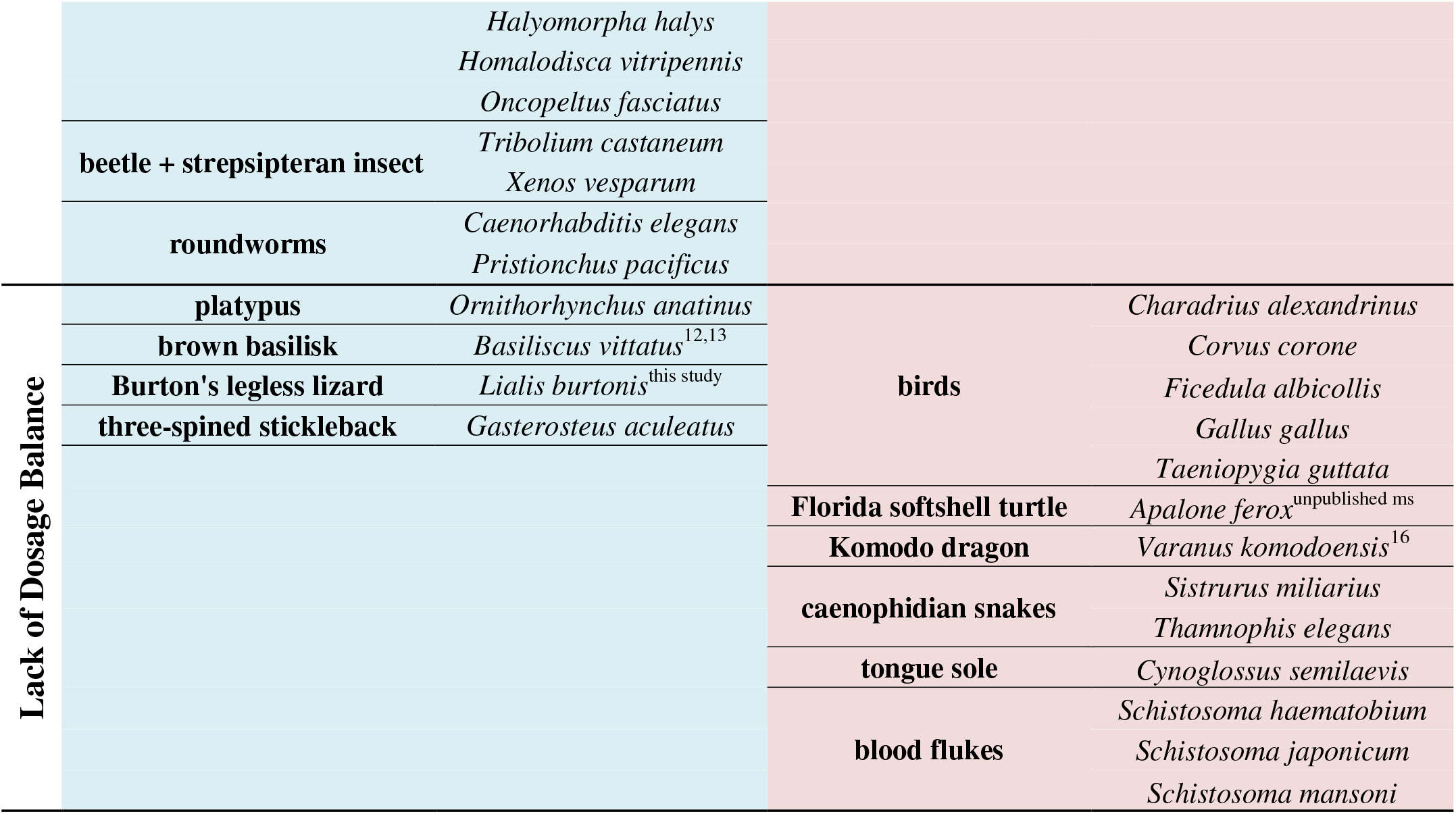
Summarization of the current knowledge on presence/absence of dosage balance across animals. Animal species are split to groups reflecting putative independent origins of sex chromosomes (see Ref. 62-65 for evidence on homology of sex chromosomes in dipteran insects). Most evidence were taken from the review by Gu and Walters^6^, supplemented by newer data (references in the individual species in the table).

Pygopodids (legless or flap-footed lizards) are a small family of 45 species of gecko lizards^28^ native to Australia and New Guinea. Pygopodids are the only lineage within the gekkotan radiation that possess an attenuate, snake-like body plan lacking limbs and digits, retaining only small flaps where rear legs would normally be^29^. Up until now, information on their sex determination is limited to largely cytogenetic evidence in four species: XX/XY sex chromosomes were reported in *Aprasia parapulchella*^30^ and *Delma butleri*^31^, and the X1X1X2X2/X1X2Y sex chromosomes in *Lialis burtonis* and *L. jicari* likely evolved via a fusion of an ancestral X with an autosome^32,33^. Male heterogamety in *L. burtonis* was confirmed by finding several male-specific anonymous molecular markers in RAD sequencing^25^. However, the homology of sex chromosomes among pygopodids and with sex chromosomes in other amniote lineages remains unknown.

In order to expand our knowledge on the evolution of sex chromosomes and gene dose regulatory mechanisms in amniotes, we tried to identify the sex chromosome gene content of the pygopodid Burton’s legless gecko (*Lialis burtonis*), where XX/XY sex determination was previously identified by cytogenetic methods. Here, we used an mRNA-seq-based pipeline to identify genes located on the X chromosome and a real-time quantitative PCR (qPCR) method to validate the candidate X-specific genes. Subsequently, the qPCR approach was further used to explore the homology of sex chromosomes among pygopodid geckos, while mRNA-seq data were used to explore the gene dose regulatory mechanism regulating the gene dose imbalance between sexes of X-specific genes in *Lialis burtonis*.

## Material and methods

### Animal sampling and DNA/RNA isolation

Tissue or blood samples were collected from both sexes of five species of pygopodids: *Aprasia parapulchella, Delma inornata, Lialis burtonis, Lialis jicari* and *Pygopus nigriceps* (Table S1). The processing of the biological material was carried out by accredited researchers and under the supervision and with the approval of the Ethics Committee of the Faculty of Science, Charles University in Prague followed by the Ministry of Education, Youth and Sports of the Czech Republic (permission 8604/2019-7).

Genomic DNA from all specimens was extracted using a DNeasy Blood and Tissue Kit (Qiagen, Valencia, CA, USA). Total RNA from the blood of two females and four males of *L. burtonis* and one male of *L. jicari* was extracted using TRIzol reagent (Invitrogen, Carlsbad, CA, USA) according to the manufacturer protocols. The quantity and purity of the extracted DNA and RNA samples were estimated using a NanoDrop ND-2000 spectrophotometer (Thermo Fisher Scientific Inc, Waltham, MA, USA).

### *RNA sequencing and identification of X-specific genes in* L. burtonis

Barcoded stranded mRNA-sequencing libraries were constructed from the total RNA samples from six individuals of *L. burtonis* and one individual of *L. jicari* by GeneCore (EMBL, Heidelberg, Germany) using the Illumina TruSeq mRNA v2 sample preparation kit (Illumina, San Diego, CA, USA) with poly-A mRNA enrichment. The libraries were pooled in equimolar amounts and loaded on the Illumina NextSeq 500 sequencer and 85 base pairs (bp) were sequenced bidirectionally. The raw Illumina reads were deposited in GenBank database under the BioProject PRJNA623146.

The raw Illumina reads were trimmed for adapters and low quality bases in Trimmomatic^34^, according to the default parameters. Reads with the size less than 50 bp were removed from the dataset, resulting in a final dataset of 40-80 million reads per specimen. Trimmed reads were checked for quality in FASTQC^35^ and MULTIQC^36^.

Trimmed reads from a single male of *L. burtonis* were assembled *de novo* with Trinity v2.8.5^37^. The assembled transcripts were compared with BLASTn to the reference transcriptomes of *Anolis carolinensis*, *Chrysemys picta*, *Gallus gallus*, *Gekko japonicus*, *Pelodiscus sinensis*, *Pogona vitticeps* and *Python molurus*. Transcript sequences of *L. burtonis* with higher than 70% similarity spanning over 150 bp of homologous sequences to a reference transcriptome were selected for further analyses, resulting in a final dataset of 64,432 annotated transcripts. The Illumina reads from all five male pygopodid specimens were independently mapped to our *L. burtonis* reference transcriptome using Geneious Prime. Consensus sequences from the assembly were exported, treating polymorphic sites (for example SNPs) in all sequences as ambiguous bases. Transcript regions with coverage below 10× and size less than 500 bp were removed from the dataset.

The Y chromosomes in both species of the genus *Lialis* contain extensive heterochromatic blocks and accumulations of repetitive motifs, indicating a high degree of degeneration of the Y chromosome^33^. Comparative genome hybridization showed that the Y and X chromosomes differ significantly in sequence content^33^. Degenerated Y and W sex chromosomes have usually lost, in their non-recombining region, most of the genes present on their X or Z counterparts, respectively. Single-copy X and Z-specific loci should contain just a single allele in the genome of the individuals from the heterogametic sex. Therefore, we can uncover candidates for such hemizygous loci based on the constant lack of SNPs in homologous transcripts from all specimens of the heterogametic sex. However, homozygous autosomal and pseudoautosomal loci might also not possess SNPs in their transcripts. To differentiate between these categories, we took advantage of the high level of conservation in chromosome synteny across sauropsids^39,40^. We assume that genuine X-specific (hemizygous) genes from male individuals should form a syntenic chromosome block enriched in loci without SNPs, but false positive (homozygous) genes should be scattered randomly across chromosomes^27^. We assigned as many transcripts of *L. burtonis* as possible to putative syntenic blocks according to chromosomal position of their orthologous genes in the chicken (*Gallus gallus*, GGA) genome (www.ncbi.nlm.nih.gov/genome). We used the chicken genome because it is well assembled and annotated compared to other avian and reptile genomes. We determined which syntenic blocks defined by chicken chromosomes are unusually enriched in loci without SNPs. Such blocks were identified as significant outliers from the linear regression between the number of genes without SNPs in a given putative syntenic block and the total number of expressed genes in a given block. For this analysis, we filtered out all transcripts with < 500 bp in length and all gene duplicates, i.e. each gene was represented by a single transcript in the dataset. Genes that lacked SNPs in all five males were considered candidate X-specific genes (i.e. those on the X chromosome but absent in the degenerated part of the Y chromosome). The differences in gene copy numbers between sexes triggered by the degeneration of the Y chromosome can also be directly measured by quantitative real-time PCR (qPCR) applied to genomic DNA^16,27,41,42^. In *L. burtonis* we used this approach for the validation of X-specificity in a subset of loci from the candidate putative syntenic blocks. Primer pairs were designed for the amplification of the 120–200 bp exon fragments of autosomal control genes and candidate X-specific genes in the Primer-BLAST software^43^ using Primer3 approach^44^. The qPCR with DNA template was carried out in a LightCycler II 480 (Roche Diagnostics, Basel, Switzerland) with all samples run in triplicates (for the list of genes and primers see Table S2). The qPCR protocol and the formula for the calculation of the relative gene dose between sexes have been presented previously^45^. A relative male-to-female gene dose ratio (r) of 0.5 is expected for X-specific genes and of 1.0 for autosomal and pseudoautosomal genes, and genes with poorly differentiated gametologs. We recently used similar methodology to discover sex-linked genes in lacertid and anguimorphan lizards and in the gecko genus *Paroedura*^16,27,42^.

### qPCR test of homology of sex chromosomes in pygopodid geckos

Candidate hemizygous genes in *L. burtonis* were tested for X-specificity in four additional pygopodid species (*Aprasia parapulchella, Delma inornata, Lialis jicari* and *Pygopus nigriceps*) using the qPCR technique (for the list of genes and primers see Table S2) to explore sex chromosome homology. The tests and calculations were performed as described above.

### Test of dosage balance in L. burtonis

We used transcriptome data from two females and four males of *L. burtonis* to test for dosage balance of the X-specific genes. FPKM (Fragments Per Kilobase Million mapped reads) expression values were independently calculated for each transcript with average read coverage > 10 across all specimens, from data provided by the Geneious Prime “map to reference” assembler. Subsequently, we computed the average sex-specific FPKMs for each transcript as the mean value from the two females and the four males, respectively. Gene expression may vary significantly between individuals of the same sex not due to gene copy number, but to physiological parameters (e.g. age, fitness, sickness, reproductive stage). Therefore, we excluded from further analysis all transcripts that had high variation among specimens of the same sex (i.e. variation more than 30% of the mean standard deviation). The duplicities in gene identity were filtered out. For the analysis, we kept only the transcript with the smallest FPKM value in males in each gene. However, the results of the following analyses led to the same conclusion even without such strict filtering of transcripts.

We applied the analysis of covariance (ANCOVA) with log10-transformed average male FPKM as the dependent variable, chicken chromosome as the factor representing the grouping of genes to putative syntenic blocks and log10-transformed average female FPKM as the covariate. We also compared ratios of average male FPKM to average female FPKM between the putative syntenic block determined as X-specific and other syntenic blocks (putative autosomes) by analysis of variance (ANOVA).

## Results

### Candidate sex chromosome genes in L. burtonis

We confidently assigned 7,718 individual genes of *L. burtonis* to chicken chromosomes (Table S3). The total number of genes linked to individual chicken chromosomes correlates tightly with the expressed genes of *Lialis burtonis* we assigned to them (Pearson’s *r* = 0.98, *n* = 34, *p* < 0.0001; Fig. S1), which demonstrates that individual chicken chromosomes are more or less uniformly represented in the pygopodid transcriptomes.

The number of *L. burtonis* genes per individual chicken chromosomes correlates well with the number of genes of *L. burtonis* without SNPs across the same chromosomes. There is only one significant outlier (the fourth largest chicken chromosome, GGA4) from this relationship. As GGA4 emerged relatively recently in the chicken ancestor via fusion of two ancestral chromosomes now largely forming small (p) and large (q) arm of GGA4^46^, we further analysed genes from GGA4p and GGA4q separately to resolve the gene content of pygopodid sex chromosomes. The residual analysis showed that only GGA4q is the significant outlier from otherwise tight relationship (*r* = 0.83, *n* = 35, *p* < 0.0001; Fig. S2) between the number of *L. burtonis* genes without SNPs versus those assigned to individual chicken chromosomes. The standard residual of GGA4q from this linear regression is very large (4.57), suggesting that this putative syntenic block is exceptionally enriched for genes without SNPs. The residuals of all the other chicken chromosomes including GGA4p are in the range between −1.40 and 1.39.

### Sex chromosome homology in pygopodid geckos

We tested four candidate X-linked genes in *L. burtonis* (*elf2, maml3, noct, rab33b*) with synteny to the q arm of chicken chromosome 4 using qPCR. The genes *cabin1*, *derl3*, *fbxo33*, *rag1*, *ubr5* and *usp12* were used as positive autosomal controls; the gene *mecom* was used for the normalization of the qPCR values (Table S2). Our qPCR experiments confirmed that the tested loci from syntenic block GGA4q are X-specific in all five tested pygopodid species (Fig. 1). Our results demonstrate that pygopodid geckos have homologous sex chromosomes, probably derived from their common ancestor.

**Figure 1:**
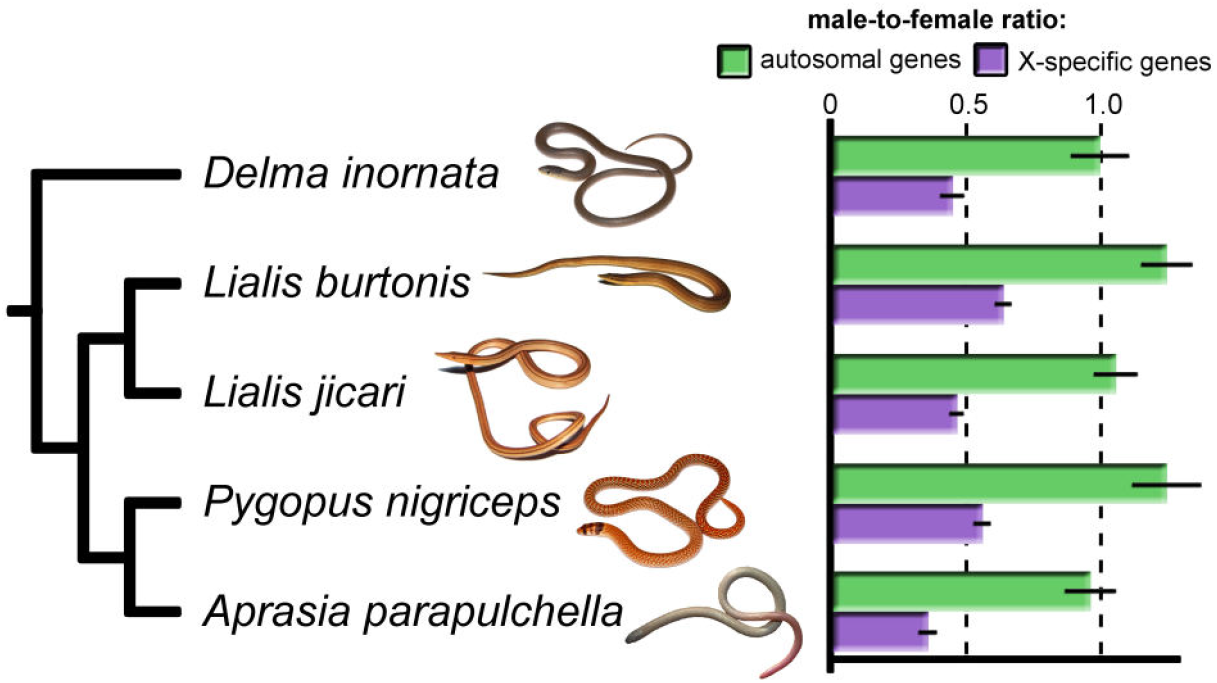
Average relative gene dose ratios between sexes for autosomal genes and X-specific genes of *Lialis burtonis* examined across five species of pygopodid lizards. The value 1.0 is expected for autosomal or pseudoautosomal genes, whereas 0.5 is consistent with X-specificity. Standard error is indicted by black bar.

### Gene dose regulatory mechanism in L. burtonis

ANCOVA showed that log-transformed average male FPKM is highly predictable by log-transformed average female FPKM (covariate: F_1_, 5057 = 203,786, *p* < 0.00001). However, at the same time, the syntenic blocks defined by chicken chromosomes strongly differ in male expression in comparison to female expression (F_29_, 5057 = 15.80, *p* < 0.00001) with chromosome GGA4q being the only very significant outlier (Fig. 2) showing that the genes linked to this syntenic block homologous to the pygopodid X chromosome are transcribed less in males. Chicken chromosomes 16 and 29-32 were represented by less than 10 genes in our *L. burtonis* dataset and were excluded from the analyses.

**Figure 2:**
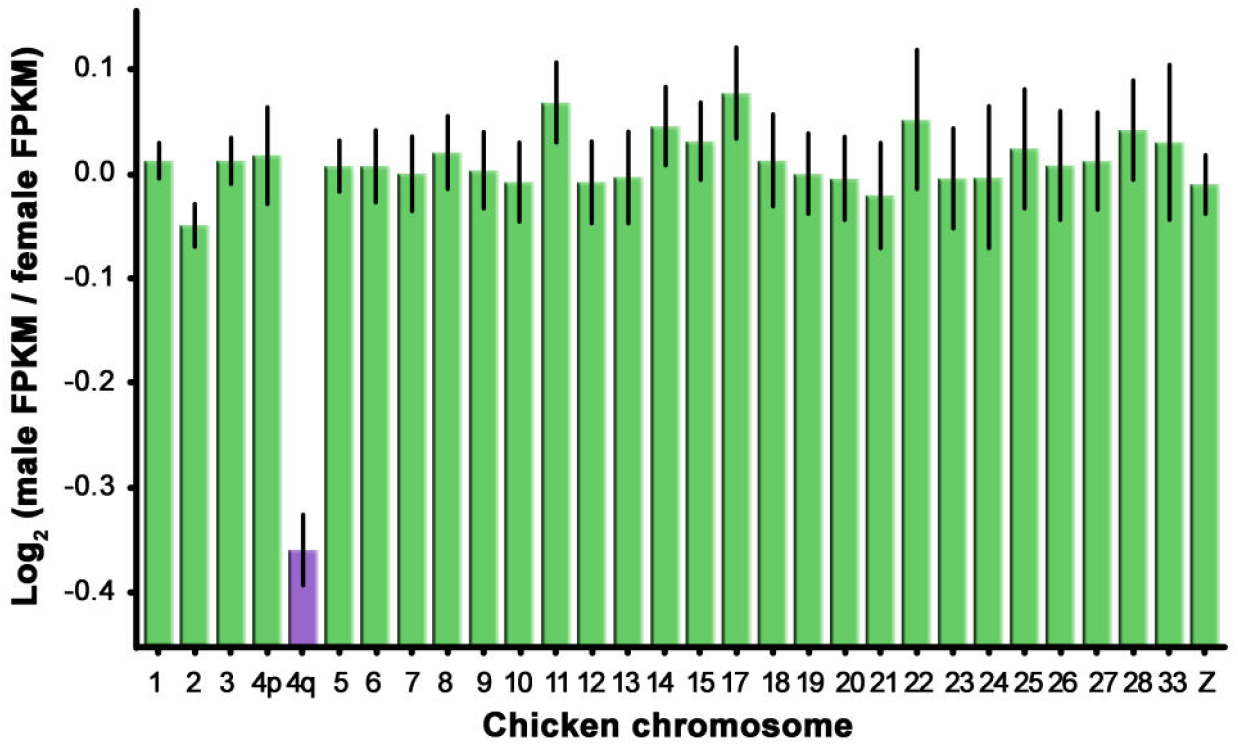
Comparison of sex-specific expression of genes from the Burton’s legless lizard among putative syntenic blocks defined by linkage of orthologs to chicken chromosomes. Note that GGA4q has exceptional sex-specific expression, which suggests that there is no dosage balance in this species.

ANOVA confirmed that the putative syntenic blocks defined by chicken chromosomes significantly differ in the log_2_-transformed ratios of average male FPKM to average female FPKM in *L. burtonis* (F_29, 5058_ = 15.80, *p* < 0.00001) and that the ratios are significantly lower in genes with orthologs linked to the chromosome GGA4q than in genes linked to other chicken chromosomes (Fig. 3).

**Figure 3:**
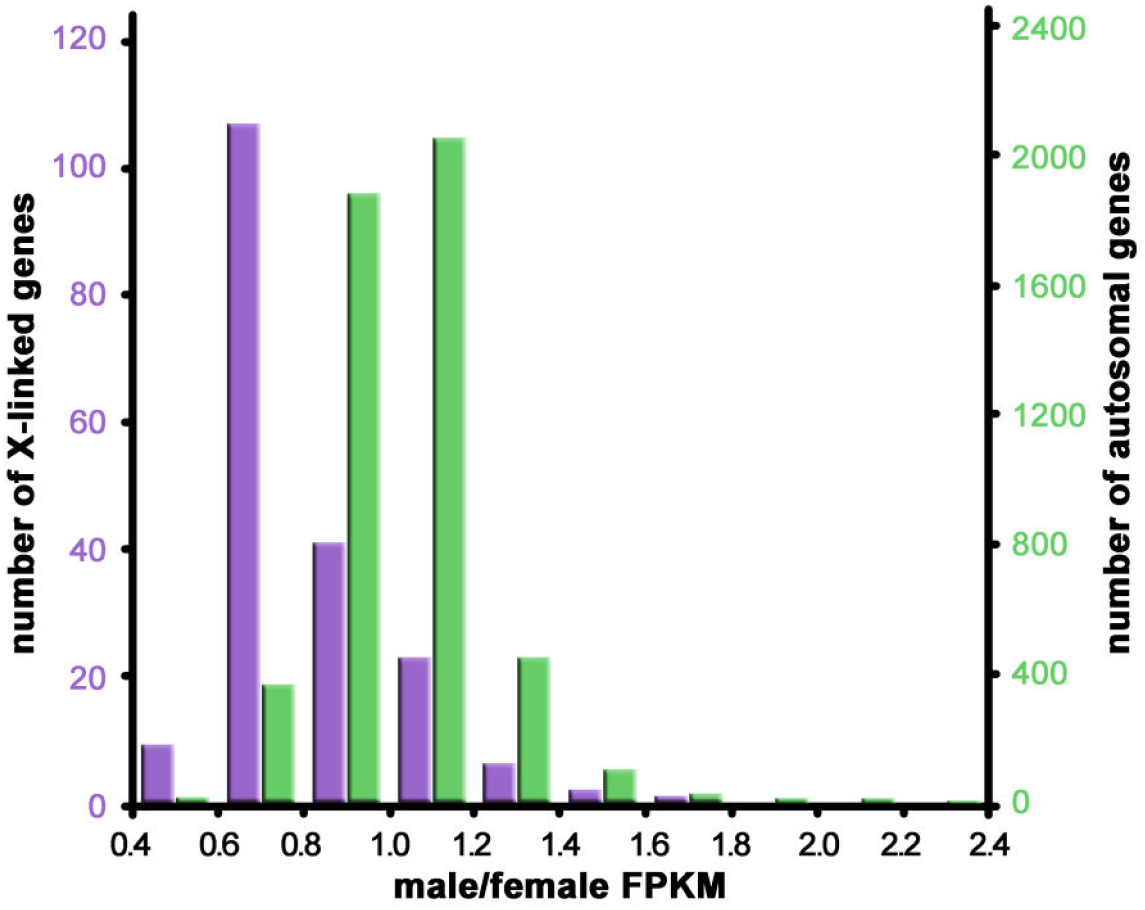
Histograms of the ratios of male to female measures of expression (FPKM) for genes linked to chicken chromosome 4q syntenic to pygopodid X chromosome and to other chromosomes in *Lialis burtonis*.

## Discussion

We identified the partial gene content of X-specific region of *L. burtonis* based on the (i) analysis of the distribution of SNPs across genes validated by the measurement of differences in gene copy numbers between sexes and (ii) by analysing expression differences of those genes between sexes. We show that the same X-specific region is shared by all sampled pygopodid species in the genera, *Aprasia, Delma, Lialis*, and *Pygopus*, despite the differences in morphology of their sex chromosomes and origin (e.g. the fusion of the ancestral sex chromosomes with an autosome leading to multiple neo-sex chromosomes in the common ancestor *of L. burtonis* and *L. jicari*)^33^. It seems that the differentiated XX/XY sex chromosomes in pygopodids are ancient and can be dated to the last common ancestor of living pygopodids, i.e. to at least 32 - 50 MY^47^. As female heterogamety is known in the sister group to pygopodids, the family Carphodactylidae^48^, the XX/XY sex chromosomes in pygopodids might be as old as 55 to 78 million years; the estimated age of when these two families split^47,49,50^. The pygopodid sex chromosomes are homologous to chromosome 4q of the chicken and the human chromosome 4. It seems that sex chromosomes in the pygopodid ancestor evolved independently from sex chromosomes of other amniotes, as no amniote group studied to date with known partial gene content of sex chromosomes share sex-linked gene content with pygopodids^27^, including three other gekkotan lineages: *Phyllodactylus wirshingi* (its ZZ/ZW chromosomes are syntenic with chicken Z; GGAZ), *Gekko hokouensis* (GGAZ as well, but likely independently derived), and the geckos of the genus *Paroedura* (GGA4p and GGA15)^27,51,52^. It should be noted that previously reported synteny of amniote sex chromosomes with chromosome GGA4 in lacertid lizards, geckos of the genus *Paroedura*, and therian mammals involved the small arm (GGA4p) not the larger arm (GGA4q) of the fourth chicken chromosome.

Genes linked to sex chromosomes in *Lialis burtonis* are expressed in blood cells significantly less in males in comparison to females (Figs. 2,3), suggesting lack of dosage balance between sexes in the expression of X-specific genes, and likely also of the global dosage compensation mechanism. Although dosage balance is lacking in all four amniote lineages with independently evolved ZZ/ZW sex chromosomes (i.e. birds, caenophidian snakes, a trionychid turtle, and the Komodo dragon), it is present in only two (i.e. eutherian mammals and the green anole) out of five studied lineages of amniotes with male heterogamety (reviewed in Table 1). This study adds Burton’s legless lizard to platypus and basilisks as another exception to the rule concerning differences in gene dose regulatory mechanisms between male and female heterogamety in amniotes.

To test whether male heterogamety is strictly linked to dosage balance in amniotes, we summarized the current state of knowledge concerning dosage balance across animals (Table 1). Dosage balance was studied in 22 lineages representing equal number of putative independent origins of sex chromosomes. In contrast to the classical models for the evolution of gene dose regulatory mechanisms, lineages with male heterogamety are not significantly more likely to possess dosage balance between sexes in the expression of genes linked to sex chromosomes than lineages with female heterogamety (Fisher’s exact test: *p* = 0.074). Moreover, the ratio can even be biased in favour of the tested hypothesis; e.g. nematodes in fact do not represent the difference in expression between males and females, but between males and hermaphrodites^53,54^. Also, we grouped species according to putative independent evolution of their sex chromosomes based on sex-linked gene content, but due to gaps in knowledge we were not able to separate independent origins of gene dose regulatory mechanisms. It is important especially in insects, which are overrepresented in the studies on gene dose regulatory mechanisms (Table 1). Most insect lineages have male heterogamety and the origin of an epigenetic mechanism ensuring dosage balance of X-linked genes could be ancient and independently co-opted for regulation of expression of sex-linked genes even after turnover of sex chromosomes. On the other hand, sex chromosomes of marsupial and placental mammals are likely homologous, but their dosage compensating mechanisms are probably not^55^. Going forward, the sampling of lineages should be increased and we should focus on the test of homology of gene dose regulatory mechanisms and sex chromosomes. However, it seems that the earlier recognized pattern of a dichotomy in gene dose regulatory mechanisms between male and female heterogamety could be the result of limited sampling instead of a systematic difference.

The important question remains what (if anything) besides male and female heterogamety determines whether a lineage would evolve global dosage balance in the expression of X- and Z-specific genes or not. We suggest that it is related to the general mechanisms of sex determination, which generally work in two ways: sex determination might be controlled either by the copy number of X or Z-linked loci per cell (i.e. gene dosage), or by a dominant W or Y locus^56^. We hypothesize that the dosage-dependent sex determination can work only in the absence of global dosage balance between sexes at least at the time of the expression of the sex-determining locus. In contrast, the sex determination based on a dominant factor on Y and W chromosomes is compatible with both presence and lack of a dosage balance influencing chromosome-wide expression of X and Z-linked genes. Unfortunately, our knowledge on the identity and function of sex determining loci together with information on gene dose regulatory mechanisms is sporadic and restricts the testing of our hypothesis, yet the limited existing information is in agreement with this hypothesis. Only lineages with sex determination controlled by the gene dose of X or Z-linked loci per cell are informative for the testing. In support of this hypothesis, both studied lineages with female heterogamety likely relying on the dosage-dependent mechanism, i.e. birds and caenophidian snakes^57,58^, have a lack of global dosage balance^10,15^. At first sight, two model organisms, the fruit fly *Drosophila melanogaster* and the nematode worm *Caenorhabditis elegans*, represent a contradictory case, since it is textbook knowledge that their sex determination primarily relies on the number of copies of the X chromosome (in correlation to autosomes ratio), but at the same time they have global dosage compensation^59,60^. However, when inspected more closely, these cases in fact do not contradict our hypothesis: dosage compensation in fruit flies and worms is triggered only later in development, and thus does not interfere with the earlier sex-determination mechanisms based on copy numbers^60,61^, which illustrates that detailed knowledge on molecular machinery and timing of particular steps will often be needed for testing mechanistic hypothesis on the evolution of gene dose regulatory mechanisms.

## Author Contributions

M.R.: experimental part and bioinformatics; L.K.: statistics; T.G., S.N., A.G., T.E.: provided part of the material and useful consultations; M.R., L.K.: conceived the project and wrote the first draft. All authors contributed to the final form of the manuscript and held responsible for its content.

## Acknowledgement

The authors would like to express their gratitude to Jana Thomayerová for technical support. M.R. and L.K. were supported by Czech Science Foundation (GAČR) project No. 19-19672S, M.R. also by Charles University PRIMUS/SCI/46 and Research Centre program (204069).

## Supplementary information

**Figure S1:**
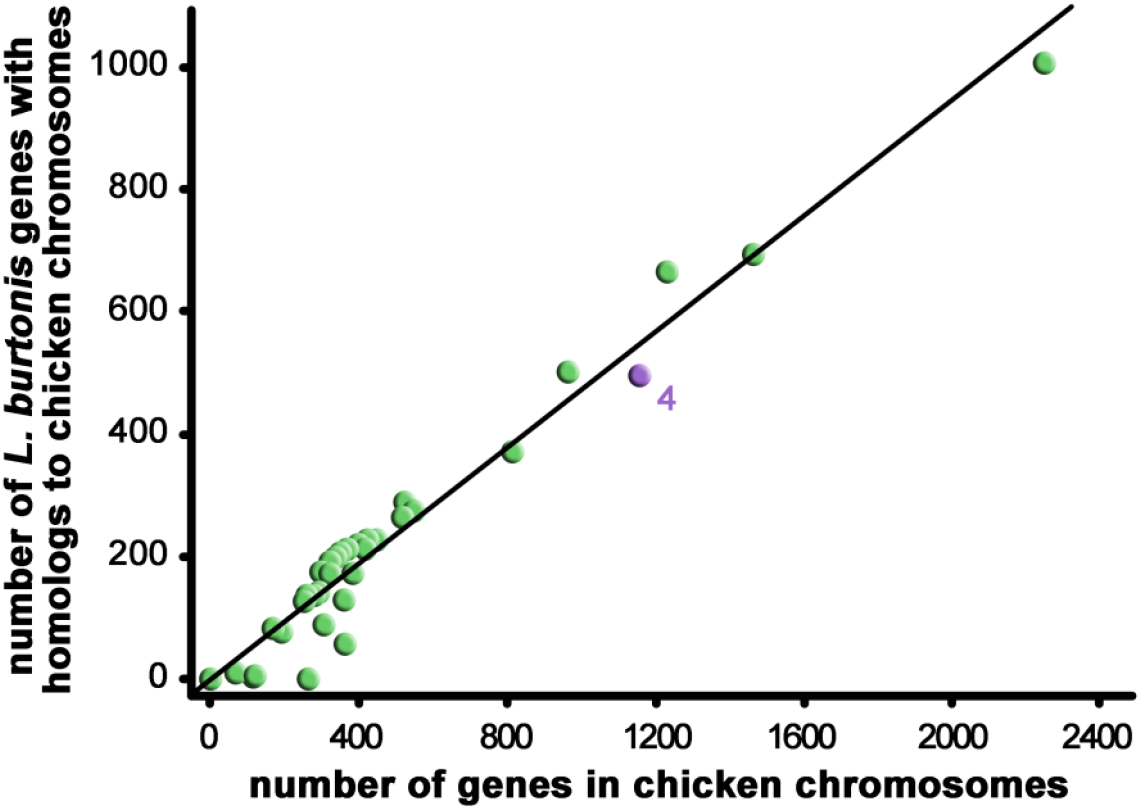
The relationship between the total number of genes linked to particular chromosomes in chicken and the number of the expressed genes in *Lialis burtonis* linked to these chromosomes. The putative syntenic block with homologs linked to GGA4 is indicated.

**Figure S2:**
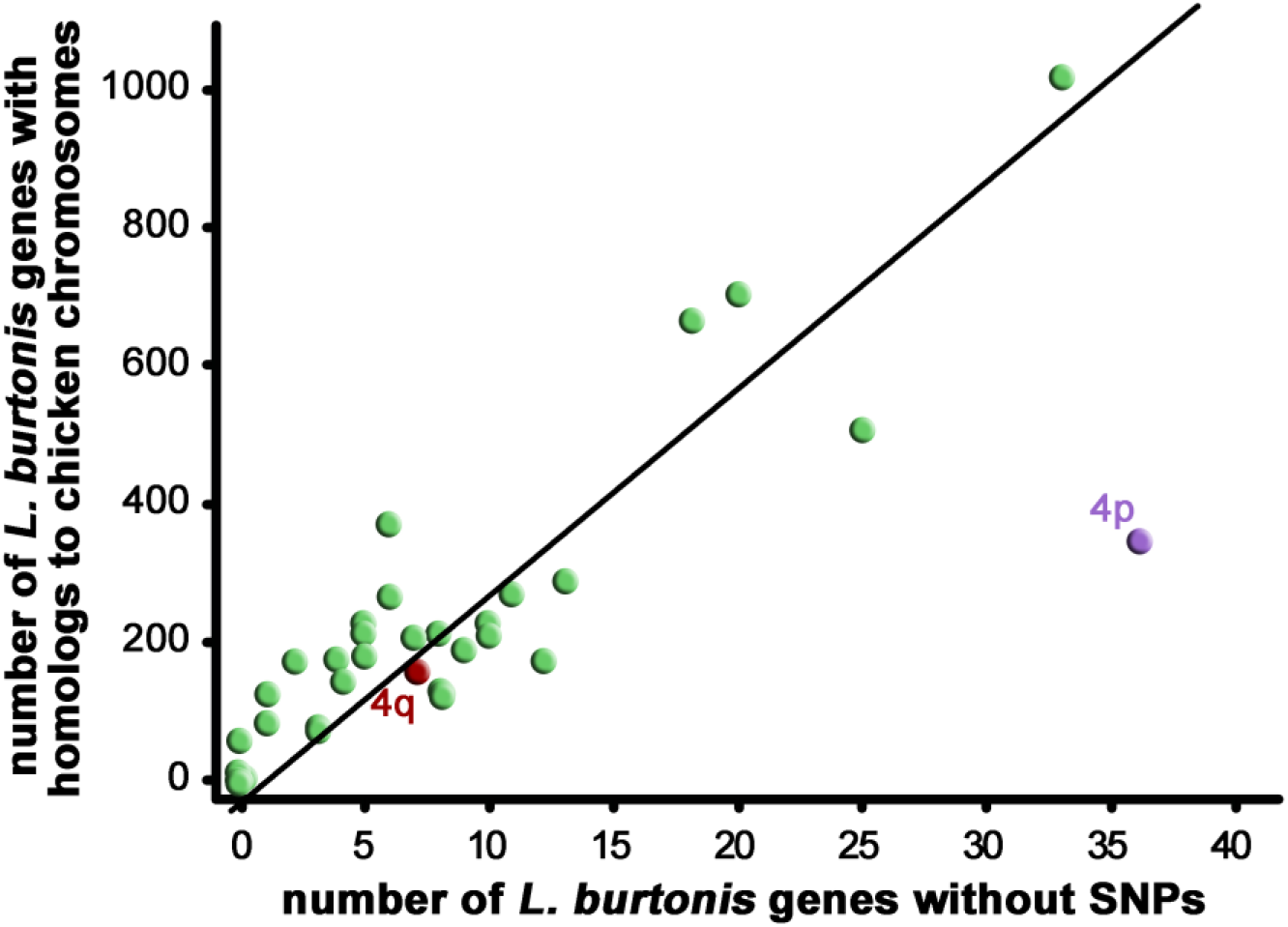
The relationship between the number of *L. burtonis* genes per individual chicken chromosomes and the number of genes of *L. burtonis* without SNPs across the same chromosomes. The significant outlier (GGA4q) from this relationship is assigned. Note that GGA4p is not outlying from the general pattern.

**Table S1:** List of pygopodid geckos, per species and sex, used in the current study.

**Table S2:** List of genes, primers and relative gene dose ratios (r) between males and females for autosomal control and X-specific genes in five pygopodid species (r = 1 corresponds to autosomal or pseudoautosomal position, while r = 0.5 corresponds to X-specificity). The Cp value for the gene mecom was used for normalization from the same run. The symbol “x” means that qPCR test was not successful (e.g. lack of amplification or presence of a secondary product).

**Table S3:** List of genes analysed from blood transcriptome, their homology in the genomes of the green anole *Anolis carolinensis*, the common wall lizard *Podarcis muralis* and the chicken *Gallus gallus* and their male to female FPKM expression value ratio.

